# Screening for insulin-independent pathways that modulate glucose homeostasis identifies androgen receptor antagonists

**DOI:** 10.1101/427831

**Authors:** Sri Teja Mullapudi, Christian S. M. Helker, Giulia L.M. Boezio, Hans-Martin Maischein, Anna M. Sokol, Johannes Graumann, Stefan Guenther, Hiroki Matsuda, Stefan Kubicek, Yu Hsuan Carol Yang, Didier Y.R. Stainier

**Author notes:** Correspondence to: Didier Stainier, Department of Developmental Genetics, Max Planck Institute for Heart and Lung Research. Ludwigstrasse 43, 61231 Bad Nauheim, Germany.

## Abstract

Pathways modulating glucose homeostasis independently of insulin would open new avenues to combat insulin resistance and diabetes. Here, we report the establishment, characterization and employment of a vertebrate ‘insulin-free’ model to identify insulin-independent modulators of glucose metabolism. *insulin* knockout zebrafish recapitulate core characteristics of diabetes and survive only up to larval stages. Utilizing a highly efficient endoderm transplant technique, we generated viable chimeric adults that provide the large numbers of *insulin* mutant larvae required for our screening platform. Using glucose as a disease-relevant readout, we screened 2233 molecules and identified 3 that consistently reduced glucose levels in *insulin* mutants. Most significantly, we uncovered an insulin-independent beneficial role for androgen receptor antagonism in hyperglycemia, mostly by reducing fasting glucose levels. Our study proposes therapeutic roles for androgen signaling in diabetes and, more broadly, offers a novel *in vivo* model for rapid screening and decoupling of insulin-dependent and -independent mechanisms.

## Introduction

Characterized by the inability to control blood glucose levels, diabetes is a metabolic disease of major socio-economic concern. Blood glucose levels are regulated by multiple tissues including the pancreas, muscle, liver, adipocytes, gut and kidney (DeFronzo, 2009). Signals from endocrine hormones are integrated at each tissue to effectively maintain glucose homeostasis, and aberrations in this interplay underlie the pathogenesis of diabetes. Currently, seven classes of antidiabetic drugs exist, of which only three function without increasing circulating insulin levels and only one that definitively functions independently of insulin (Chaudhury et al., 2017). Restoring normoglycemia independently of insulin secretion or action could delay disease progression as an improved glycemic status can restore β-cell mass and function (Wang et al., 2014). Lower dependence on insulin-stimulating therapies can also prevent hyperinsulinemia-driven insulin resistance (Shanik et al., 2008) and obesity (Mehran et al., 2012). In contrast to insulin stimulators, Biguanides (E.g.: Metformin) and Thiazolidinediones (E.g.: Pioglitazone) are effective antidiabetic agents that primarily sensitize tissues to insulin (reviewed by (Soccio et al., 2014; Rena et al., 2017)). Likewise, sodium-glucose transporter 2 inhibitors (E.g.: Dapagliflozin) have a complementary mechanism of reducing glucose reabsorption in the kidney (Bailey et al., 2013). Increasing evidence points to additional molecular pathways that can improve metabolic homeostasis independently of insulin, for instance, using leptin therapy (Neumann et al., 2016) or during exercise (Stanford & Goodyear, 2014). Interestingly, currently prescribed drugs were discovered from their historical use in herbal medicine (Ehrenkranz et al., 2004; Bailey, 2017) or from screens directed against hyperlipidemia (Fujita et al., 1983). However, so far, an unbiased search for insulin-independent pathways controlling glucose homeostasis has remained elusive, primarily due to the lack of a disease-relevant animal model for rapid screening. Due to its high fecundity and amenability to chemical screening, the zebrafish serves as an excellent platform to study diabetes, and it has been successfully used to study β-cell mass and activity, as well as glucose metabolism (Andersson et al., 2012; Gut et al., 2013; Tsuji et al., 2014; Nath et al., 2015; Li et al., 2016; White et al., 2016; Matsuda et al., 2018). Here, using the zebrafish model, we generated an innovative drug discovery strategy, screened chemical libraries and specifically identified insulin-independent effects of androgen signaling on glucose homeostasis.

## Results and Discussion

### *insulin* is crucial for zebrafish metabolic homeostasis

Insulin plays a central role in glucose homeostasis by increasing glucose uptake in peripheral tissues, promoting glycogenesis in the liver and decreasing glucose production by inhibiting glucagon secretion (Aronoff et al., 2004). We generated zebrafish devoid of insulin signaling and determined the degree to which these mutants recapitulate core features of diabetic metabolism observed in mammals. The zebrafish genome contains two insulin genes - *insulin (ins)* and *insulinb (insb)*. Using CRISPR/Cas9 mutagenesis, we generated a 16 bp deletion allele for *ins* (Figure 1A) and a 10 bp insertion allele for *insb*. Although *ins* and *insb* mutant embryos appear morphologically unaffected (Figure 1 - figure supplement 1A), Insulin was entirely absent in pancreatic islets from *ins* mutants (Figure 1B), whereas, there was no observable change in *insb* mutant islets (Figure 1 - figure supplement 1B, 1C). Deletion of *ins* led to a drastic increase in total glucose levels (up to 10-fold), measured from 1 to 6 days post fertilization (dpf) (Figure 1C). Additionally, staining for lipid content using Nile Red revealed large unused yolk reserves (Figure 1D), suggesting defects in lipid absorption and processing. Due to a combination of these metabolic defects, *ins* mutants do not survive beyond 12 dpf (Figure 1E). Moreover, although 3-month old (adult) *ins +/-* animals are normoglycemic (Figure 1 - figure supplement 1D), 50 dpf (juvenile) *ins +/-* animals are noticeably smaller (Figure 1 - figure supplement 1E), consistent with a role for Insulin in growth control (Nakae et al., 2001). *insb* mutants, on the other hand, are viable and fertile. As *insb* expression is negligible beyond 48 hpf (Papasani et al., 2006; White et al., 2017) (Figure 1 - figure supplement 1F), we overexpressed *insb* under the *ins* promoter to assess if *insb* was functional. Under the hyperglycemic conditions resulting upon morpholino-mediated *ins* knockdown, *insb* overexpression successfully lowered glucose levels, thus indicating that *insb* is functional (Figure 1 - figure supplement 1G). However, due to the post-embryonic expression of *ins*, survival and metabolic homeostasis in zebrafish depends primarily on *ins*. This predominant role of *ins* distinguishes the zebrafish insulins from the redundant metabolic roles of mouse *Ins1* and *Ins2* (Duvillie et al., 1997). To further explore the nature of metabolic defects in zebrafish *ins* mutants, we probed the proteome of 108 hpf *ins* mutants and compared it to seven other proteomes from diabetic tissues of murine or human origin (Figure 1F) (Hwang et al., 2010; Mullen & Ohlendieck, 2010; Giebelstein et al., 2012; Valle et al., 2012; S.-J. Kim et al., 2014; Capuani et al., 2015; Braga et al., 2016; Zabielski et al., 2016). Strikingly, pathways like gluconeogenesis, mitochondrial dysfunction, Sirtuin signaling, and oxidative phosphorylation, which were affected in diabetic conditions across these studies, were also similarly disrupted in zebrafish *ins* mutants (Supplementary file 1). Together, these findings indicate that zebrafish *ins* is crucial for metabolic homeostasis and survival and that its absence recapitulates core features of diabetic metabolism already at larval stages.

**Figure 1.**
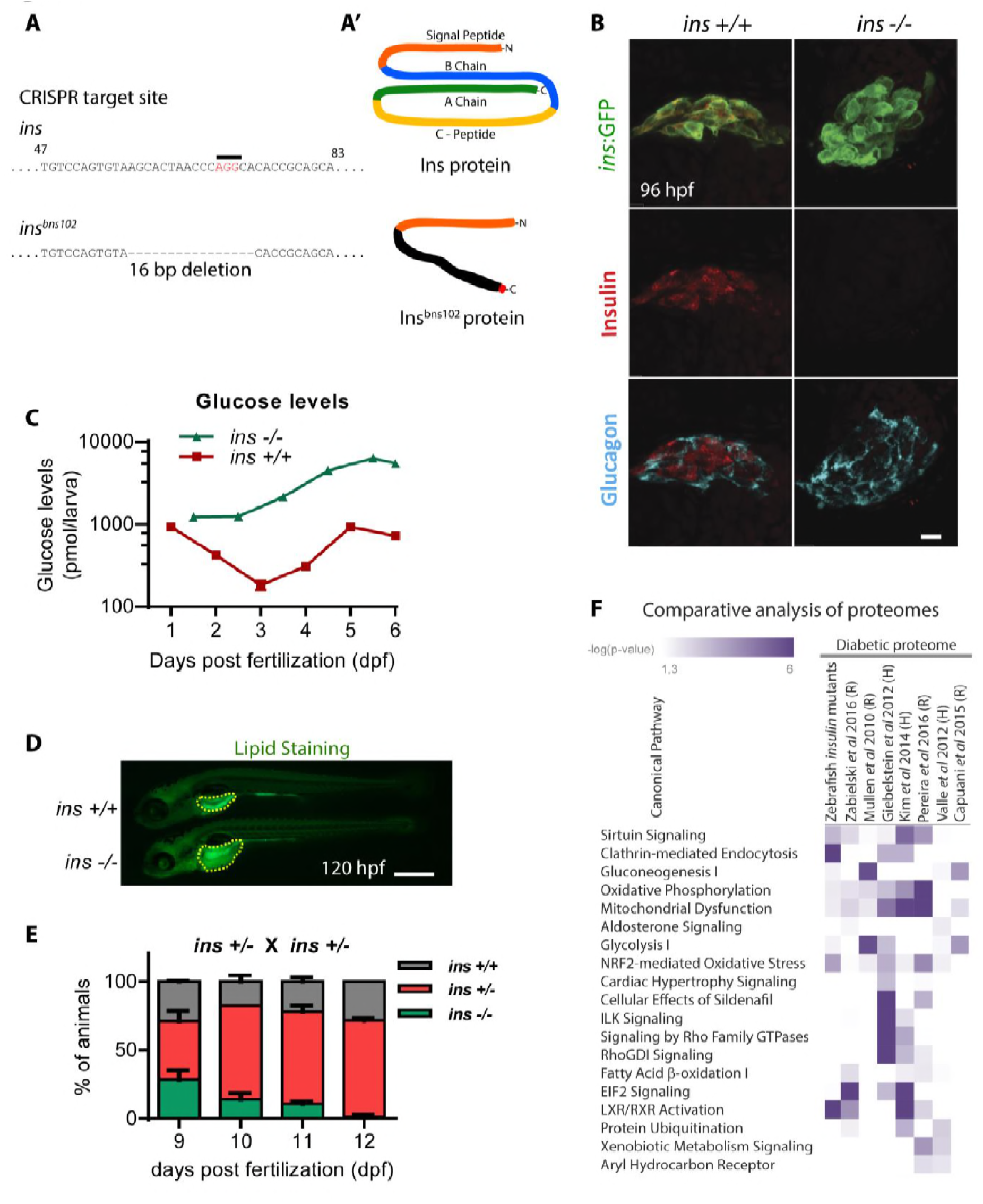
*insulin* is crucial for zebrafish metabolic homeostasis. A) CRISPR target site in the *insulin* gene, with PAM sequence highlighted, and the resulting 16 bp deletion allele (below). **A’**) Schematic of wild-type Insulin protein and the predicted mutant protein which contains novel sequence (black) and a premature stop (red). **B)** Confocal projection images of pancreatic islets in 96 hpf *Tg(ins:GFP) ins +/+* and *ins -/-* animals immunostained for Insulin (red), Glucagon (cyan). **C)** Free glucose levels measured in wild-type and mutant animals from 1 to 6 dpf; mean ± SEM, n=2-4 replicates. **D)** Nile Red staining (green) for neutral lipids in 120 hpf wild-type (top) and mutant (bottom) larvae, with yolk lipid content outlined (yellow dots). **E)** Genotype distribution from *ins +/-* incross, calculated as the percentage of total animals at each stage; mean ± SEM, n = 32 animals. **F)** Heat map of the proteomic signature of zebrafish *ins* mutants at 120 hpf compared to signatures from rodent (R) and human (H) diabetic proteome studies. Canonical pathways implicated in most studies are listed first. P-value cut-off set at < 0.05. Scale bars: 10 μm (B), 500 μm (D).

### Highly efficient endoderm transplant technique rescues *ins* mutants to adulthood

Screening of small molecules in *ins* mutants requires large numbers of mutant embryos. However, the early lethality of *ins* mutants did not allow the generation of adult animals that could be incrossed. To overcome this obstacle, we used an efficient endoderm induction (Kikuchi et al., 2001) and transplantation technique (Stafford et al., 2006) (Figure 2A) to selectively modify endodermal tissues without altering the germline. *ins +/+; Tg(ins:DsRed)* embryos were injected with *sox32* mRNA at the one-cell stage, conferring an endodermal fate on all cells. Between 3 to 4 hpf, cells were transplanted from these embryos to the mesendoderm of similarly staged embryos from *ins* +/-; *Tg(ins:RasGFP)* incrosses. This transplantation procedure was remarkably efficient at contributing to host endoderm (Figure 2 - figure supplement 1C-C’’), and nearly every transplanted embryo contained pancreatic islets with both transplanted *ins +/+* and host β-cells (Figure 2B-B’’). These chimeric animals were grown to adulthood and genotyping (Figure 2 - figure supplement 1A-B) revealed a near Mendelian ratio of mutant animals (Figure 2C). These chimeric mutant animals contain *ins +/+* endodermal tissues but retain an *ins -/-* germline, and allowed an all-mutant progeny to be obtained by incrossing.

**Figure 2.**
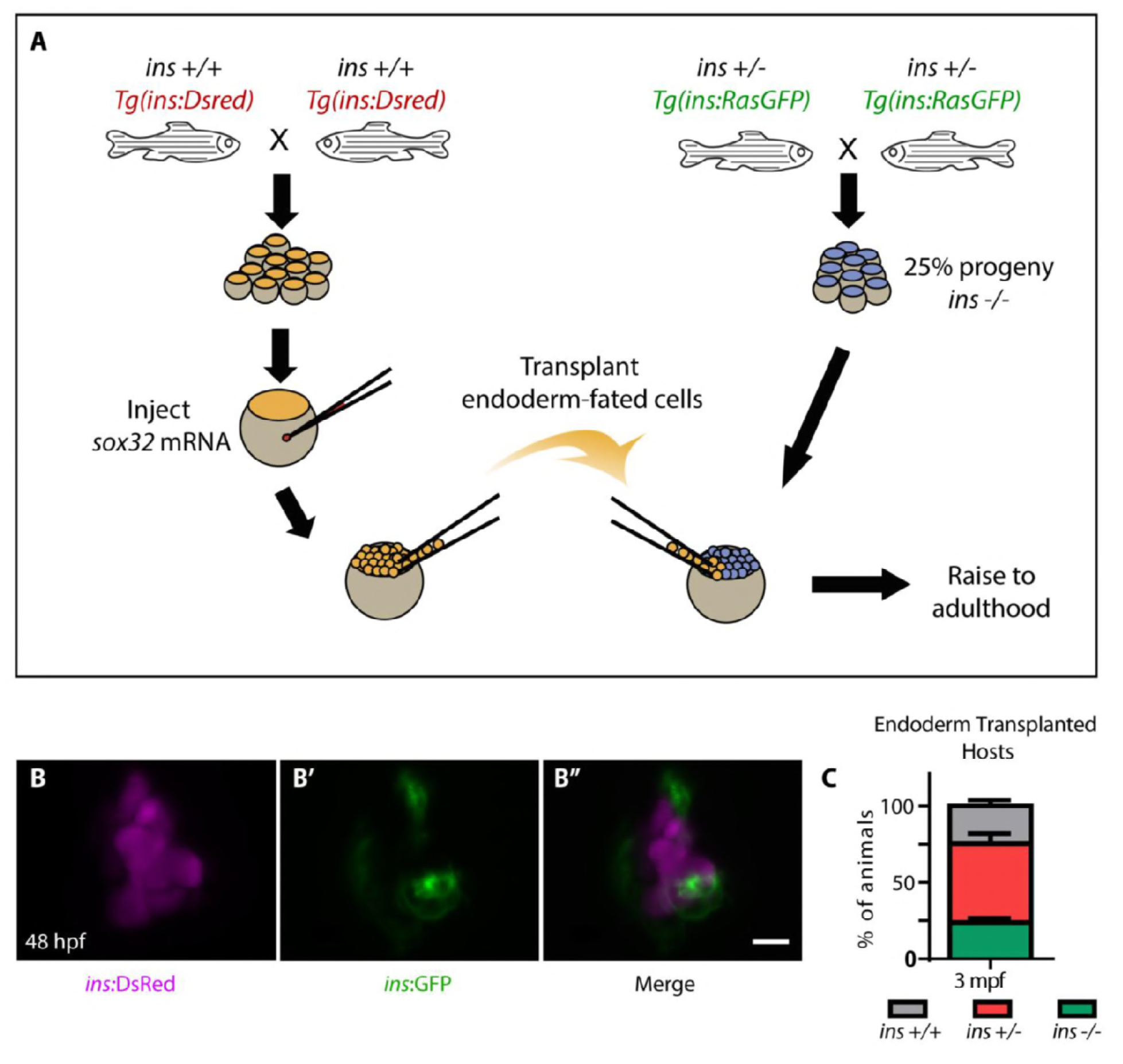
Highly efficient endoderm transplant technique rescues *ins* mutants to adulthood. A) Schematic depicting the endoderm transplantation protocol; sox32-injected *ins +/+* donor cells (orange) were transplanted into host embryos (blue) to form chimeric animals. B-B’’) Confocal projection images of pancreatic islets from 48 hpf chimeric animals show β-cells from the host (green, B’) and the transplanted *ins +/+* cells (magenta, **B)**. **C)** Quantification of genotype abundance in the raised 3 mpf chimeric animals, determined by genotyping fin tissue; mean ± SEM, n = 3 transplant experiments, 18 - 32 animals per experiment. Scale bar: 10 μm.

### Small molecule screen in *ins* mutants reveals insulin-independent modulators of glucose metabolism

With the ability to obtain large numbers of *ins* mutant embryos, we next aimed to analyze the effect of known glucose homeostasis modulators and also to screen for novel ones. We tested the effects of molecules that have been proposed to exert insulin-independent modulation of glucose homeostasis, amongst other effects. Anti-diabetics such as Metformin, Pioglitazone and Dapagliflozin, as well as the Lyn kinase activator MLR1023 (Saporito et al., 2012), were tested. We also tested Fraxidin, identified in a screen for molecules that increase glucose uptake in zebrafish (Lee et al., 2013). Surprisingly, Metformin and MLR1023 exhibited no glucose-lowering effect in *ins* mutants suggesting that they act more as sensitizers of insulin signaling rather than acting independently of insulin. On the other hand, Pioglitazone, Dapagliflozin and Fraxidin reduced glucose levels by 11, 12 and 5% respectively (Figure 3A), thus attributing part of their glucose lowering effect to an insulin-independent mechanism. Based on these data, we decided to screen chemical libraries using our model to identify molecules that could reduce glucose levels by more than 10%. To rapidly measure glucose levels in a 96-well plate format, we adapted a glucose measuring kit to be sensitive to endogenous changes in larval glucose levels (Figure 3 - figure supplement 1A-C), and established a screening pipeline (Figure 3B). We screened 2233 molecules in 2 replicates at 10 μM concentration and found 3 hits (Figure 3C) that reproducibly reduced glucose levels upon retesting with independent chemical stocks and the unmodified standard glucose measurement kit. These 3 hits - Flutamide (androgen receptor antagonist), ODQ (soluble guanylyl cyclase inhibitor (Boulton et al., 1995)) and Vorinostat (broad HDAC inhibitor (Finnin et al., 1999)) were found upon retesting multiple times to reduce glucose levels by 40, 22 and 19% respectively (Figure 3D, Figure 3 - figure supplement 1D).

**Figure 3.**
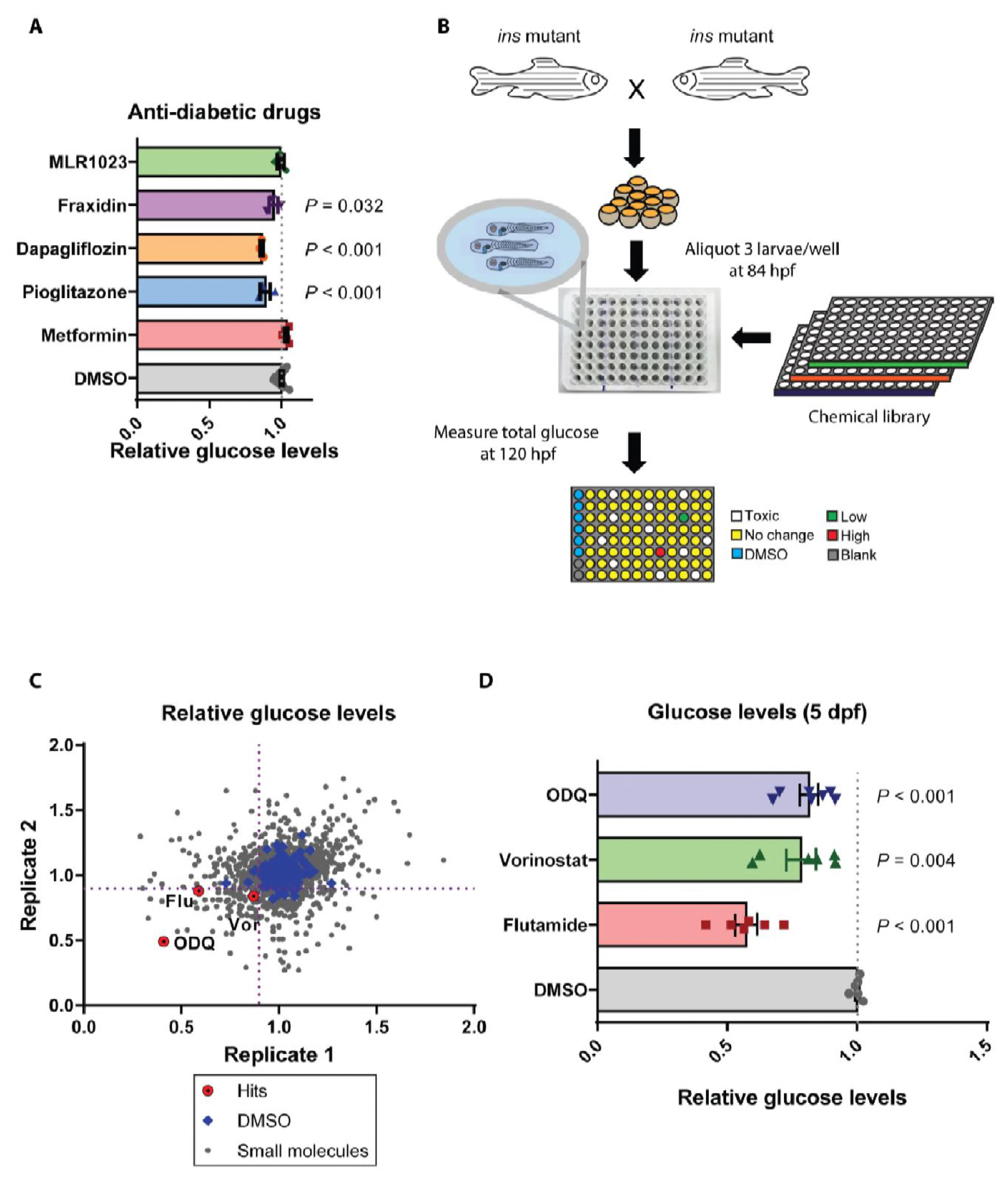
Small molecule screen in *ins* mutants reveals insulin-independent modulators of glucose metabolism. **A)** Relative glucose levels in 120 hpf *ins* mutant larvae after 36 hrs of treatment with anti-diabetic drugs (250μM Metformin, Pioglitazone, Dapagliflozin) or reported insulin mimetics (MLR1023, Fraxidin), mean ± SEM, n = 3-7 replicates. **B)** Schematic representation of the screening pipeline: *ins* mutant larvae were treated with small molecules starting from 84 hpf and total glucose levels measured at 120 hpf. **C)** Scatter-plot showing relative change in glucose levels upon treatment with 2233 small molecules. X and Y axes represent two replicates performed for each drug, with the dotted purple lines marking 0.9 on each axis. **D)** Relative glucose levels at 120 hpf upon treatment of *ins* mutants with the 3 hits - Flutamide, Vorinostat and ODQ, mean ± SEM, n = 6-7 replicates.

### Androgen Receptor (AR) antagonism regulates glucose homeostasis in hyperglycemic larval and adult animals

Given the strong reduction in glucose levels observed after Flutamide treatments, we further tested the hypothesis that glucose levels in *ins* mutants were being reduced through androgen receptor antagonism. First, Flutamide caused a dose-dependent decrease in glucose levels in *ins* mutants (Figure 4 - figure supplement 1A). Second, we treated *ins* mutants with AR antagonists of two types: (i) steroidal (Cyproterone) and (ii) non-steroidal (Nilutamide, Hydroxyflutamide, Bicalutamide, Enzalutamide). We observed consistent decrease in glucose levels across all treatments, albeit at varying efficiency (Figure 4A), possibly reflecting the specificity of these antagonists towards zebrafish AR (Raynaud et al., 1979; Teutsch et al., 1994; Tran et al., 2009). Finally, to modulate AR protein levels, we injected 1 ng of control or *ar* morpholino (MO) in one-cell stage embryos and observed a reduction of glucose levels in *ar* MO injected *ins* mutants (Figure 4B) but not in *ar* MO injected wild-type animals (Figure 4C). These data support a role for antagonizing AR specifically in hyperglycemic conditions.

**Figure 4.**
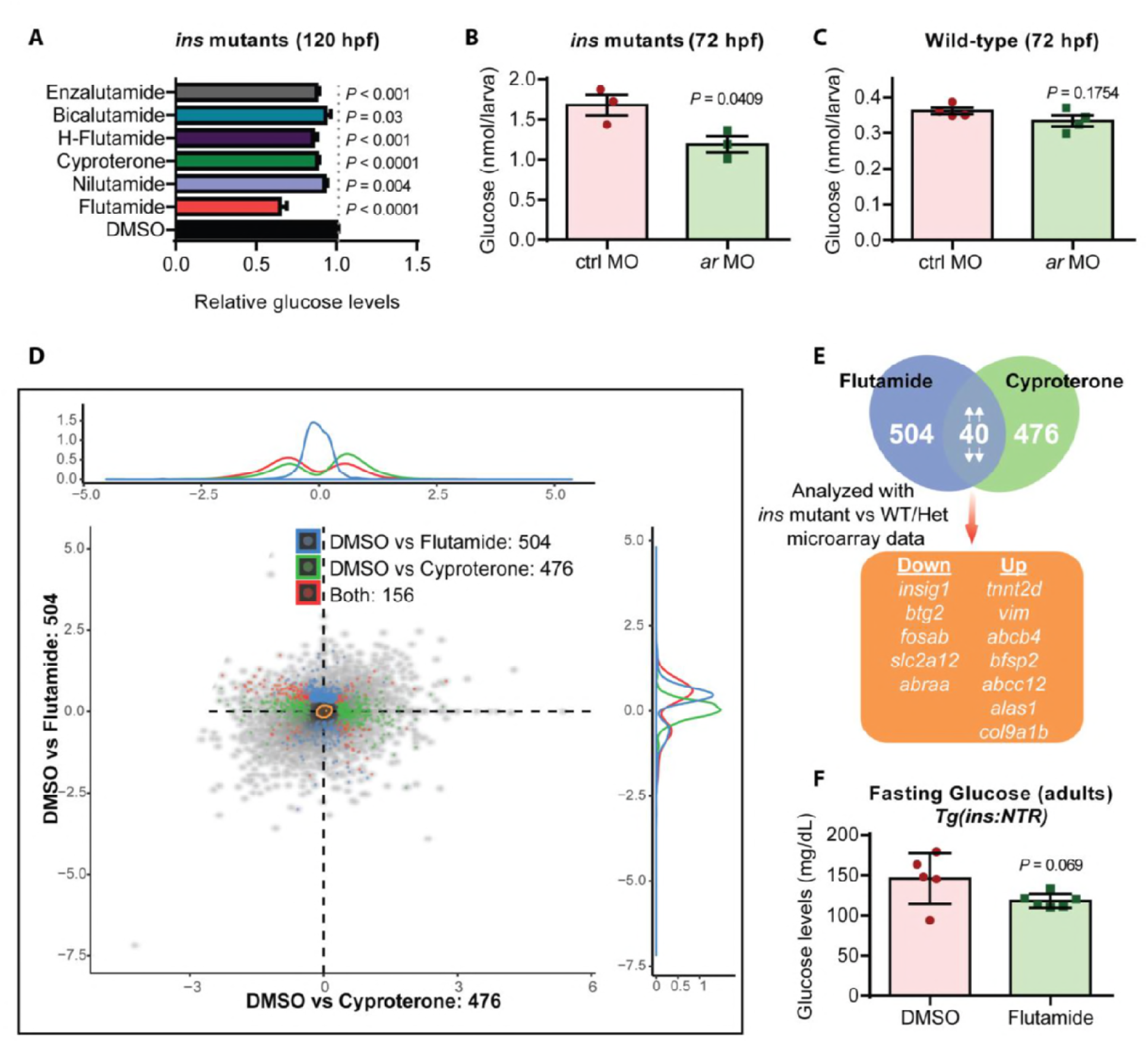
Androgen Receptor (AR) antagonism improves glucose homeostasis in hyperglycemic larvae and adults. **A)** Relative glucose levels in *ins* mutants at 120 hpf upon treatment with various AR antagonists, mean ± SEM, n = 3-7 replicates. Enzalutamide and Bicalutamide treatments were performed at 20 μM concentration. **B)** Glucose levels in 72 hpf *ins* mutants after injection with 1 ng of ctrl or *ar* MO, mean ± SEM, n = 3 replicates. **C)** Glucose levels in 72 hpf wild types after injection with 1 ng of ctrl or *ar* MO, mean ± SEM, n = 4 replicates. **D)** RNA-seq analysis of *ins* mutant larvae treated with Flutamide or Cyproterone, showing differentially expressed genes (DEGs) compared to DMSO-treated larvae in blue and green, respectively. The red dots indicate DEGs common to both treatments. **E)** Workflow used for filtering candidate genes: 40 DEGs modulated in the same direction (both up or both down) were analyzed in relation to the microarray dataset (mutant vs phenotypically wild-type 108 hpf larvae). **F)** Glucose levels measured in adult *Tg(ins:NTR)* animals after J3-cell ablation and intraperitoneal injection with vehicle (DMSO) or Flutamide; mean ± SEM, n = 5-6 animals.

A number of mechanisms have been proposed to explain the predisposition of women with androgen excess to diabetes, including insulin resistance, visceral adiposity and β-cell dysfunction (Navarro et al., 2015). Under high fat diet, a combination of neuronal and pancreatic β-cell specific roles for AR have been proposed to predispose female mice with androgen excess to diabetes (Navarro et al., 2018). Supporting this role, *ar* gene expression was observed in the zebrafish central nervous Figure system and, additionally, in the liver (Gorelick et al., 2008) (Figure 4 - figure supplement 1C). To investigate how AR antagonism mediates glucose level reduction in *ins* mutants, we evaluated the effects of antagonist treatment using transcriptomic studies. RNA-Seq analyses on 120 hpf *ins* mutants treated with Flutamide or Cyproterone revealed 504 and 476 differentially expressed genes (DEGs) compared to vehicle treated mutants (Figure 4D), respectively. Of these DEGs, 40 were regulated in parallel (both up or both down) for both AR antagonists tested, likely highlighting the common AR-specific effects. Cross-referencing these candidates with a transcriptomic comparison of *ins* mutants to phenotypically wild-type animals, led to 12 genes (Figure 4E) that were differentially expressed upon loss of *ins*, and were partially or fully restored to wild-type levels upon treatment with AR antagonists (Figure 4 - figure supplement 1B). Genes such as *btg2* and *insig1* have been reported to play crucial roles in controlling liver gluconeogenesis (Carobbio et al., 2013; Y. D. Kim et al., 2014), and they also contain two androgen response elements (AREs) close to their transcription start site (Figure 4 - figure supplement 1D). Additionally, upon intraperitoneal injections of Flutamide in hyperglycemic adult animals (Figure 4 - figure supplement 1E), we observed up to 19% lower fasting plasma glucose levels were observed (Figure 4F), likely due to reduced hepatic glucose production. Our findings corroborate the observations of better anthropometric indices previously observed with Flutamide (Sahin et al., 2004) or Metformin + Flutamide combination therapies (Gambineri et al., 2004; Amiri et al., 2014) and attribute a part of this beneficial effect to Flutamide’s insulin-independent action through AR antagonism.

In conclusion, ours is the first study reporting the generation and use of a rapid screening strategy to identify insulin-independent pathways modulating metabolism in vertebrates. Given the recent success of SGLT2 inhibitors as combination therapy in diabetes (Bailey et al., 2013), our study is an important step towards identifying more insulin-independent mechanisms governing glucose homeostasis. One of the limitations in our screen is the relatively low size of the chemical library screened. However, as the endoderm transplant technique reported here can be combined with several genetic or metabolic readouts, future studies with larger chemical libraries should unveil mechanisms governing other disease-relevant phenomena as well. Such comprehensive insight into insulin-independent mechanisms and their interactions with insulin signaling in homeostasis and disease will open new avenues for targeting therapies to treat metabolic disorders.

## Materials and Methods

### Zebrafish lines

Zebrafish husbandry was performed under standard conditions in accordance with institutional (MPG) and national ethical and animal welfare guidelines. The transgenic and mutant lines used in the study are *Tg(ins:DsRed)^m1018^* (Anderson et al., 2009), *Tg(-4.0ins:GFP)^zf5^* (Huang et al., 2001), *Tg(sox17:GFP)^s870^* (Sakaguchi et al., 2006), *Tg(ins:EGFP-HRas, cryaa:mCherry)^bns294^, Tg(ins:Flag-NTR, cryaa:mCherry)^s950^* (Andersson et al., 2012), *Tg(ins:TagRFPt-P2A-insB)*^bns285^*, ins^bns102^ (ins* mutants), and *insb^bns295^ (insb* mutants). 1, 2, 3, 4, 5 and 6 days post fertilization (dpf) correspond to 24, 48, 72, 96, 120, and 144 hours post fertilization (hpf).

### CRISPR/Cas9 mutagenesis

CRISPR design platform (http://crispr.mit.edu) was used to design sgRNAs against *ins* (targeting sequence: TCCAGTGTAAGCACTAACCCAGG) and *insb* (targeting sequence: GGATCGCAGTCTTCTCC) genes and constructs were assembled as described previously (Jao et al., 2013; Varshney et al., 2015). Briefly, a mixture of 25 pg gRNA with 300 pg Cas9 mRNA was injected into 1-cell stage wild-type embryos. High-resolution melt analysis (HRMA) (Eco- Illumina) was used to determine efficiency of sgRNAs and genotype animals with *ins* primers 5’- GTGCTCTGTTGGTCCTGTTGG-3’ and 5’-CATCGACCAGATGAGATCCACAC-3’, and *insb* primers: accordingly, 5’-AGTATTAATCCTGCTGCTGGCG-3’and 5’-GTGTAGAAGAAACCTCTAGGC-3’.

### Immunostaining, Nile Red staining and imaging

Immunostaining and imaging was performed as described previously (Yang et al., 2018). Briefly, zebrafish larvae were euthanized with incubation on ice and fixed overnight at 4°C with 4% PFA (dissolved in buffer with composition: 22.6 mM NaH_2_PO_4_, 77 mM Na_2_HPO_4_, 120 μM CaCl_2_, 117 mM sucrose, pH 7.35). After two PBS washes, the larvae were deskinned, and permeabilized using PBS containing 0.5% TritonX-100 and 1% DMSO for 1 hr. Larvae were then incubated in blocking buffer (Dako) containing 5% goat serum for 2 hrs, and incubated with primary antibody overnight at 4°C. Next, samples were washed 3 × 10 min with PBS containing 0.1% TritonX-100, incubated overnight at 4°C with secondary antibody and DAPI (10 μg/ml), washed 3 × 10 min and mounted in agarose. Antibody dilutions used are as follows: guinea pig anti-Insulin polyclonal (1:100, Thermo), mouse anti-Glucagon (1:300, Sigma), chicken anti-GFP (1:300, Aves), goat anti-guinea pig AlexaFluor568 (1:300, Thermo), goat anti-mouse AlexaFluor647 (1:300, Thermo), goat anti-chicken AlexaFluor488 (1:500, Thermo). Zeiss LSM700 (10X) and LSM800 (25X) was used to acquire data, and Imaris (Bitplane) was used for analyzing data and creating maximum intensity projection images.

Neutral lipid staining using Nile Red dye was performed at a working concentration of 0.5 μg/mL for 30 minutes in the dark, followed by acquisition of fluorescent images using an LP490 filter on a Nikon SMZ25 stereomicroscope.

### Morpholino and intraperitoneal injections

For knockdown of gene expression, the following splice-blocking antisense morpholinos (Gene Tools, LLC) were injected into one-cell embryos at the indicated dosage per embryo:

*insa* MO (4 ng, 5’-CCTCTACTTGACTTTCTTACCCAGA-3’) (Ye et al., 2016)

*ar* MO (1 ng, 5’-AGCAGAGCCGCCTCTTACCTGCCAT-3’) (Peal et al., 2011)

Standard control MO (5’ -CCT CTT ACCT CAGTT ACAATTT ATA-3’)

Intraperitoneal injections and glucose measurement in 6 month-old adult zebrafish was performed as described previously (Moss et al., 2009; Eames et al., 2010). Briefly, ablation of P- cells in *Tg(ins:NTR)* animals was performed by injecting 0.25 gm MTZ/kg body weight twice - on Day 0 and Day 1 - injecting twice improved the consistency of ablation. Flutamide (10 mg/kg) or vehicle (DMSO) was injected on Day 2, 3 and 4. For injections, animals were anaesthetized using 0.02% Tricaine. On Day 4, animals were euthanized and blood glucose was measured using FreeStyle Freedom Lite Glucose Meter (Abbott). Non-parametric Student’s t-test was used for statistical analyses and *P* values are shown in the figures.

### Larval glucose measurement

Glucose measurements were performed as described previously (Jurczyk et al., 2011), with minor modifications. After desired treatment conditions, pools of 10 animals were collected in 1.5 mL Eppendorf tubes, water removed, and frozen at -80 °C. For analysis, pools of wild-type embryos were resuspended in PBS. Samples were homogenized using a tissue homogenizer (Bullet Blender Gold, Next Advance). Glucose Assay Kit (CBA086, Merck) was used for glucose detection. Different volumes were used for resuspension and glucose detection between wild types and *ins* mutants: wild-type samples were resuspended in volume corresponding to 8 μl/animal and 8 μl was used for the glucose detection reaction. *ins* mutant embryos, due to their higher glucose content, were resuspended in volumes corresponding to 16 μl/animal and only 2 μl was used for glucose detection. Non-parametric Student’s t-test was used for statistical analyses and *P* values are shown in the figures.

### Transplantations

For the endoderm transplant experiment, *sox32* mRNA was transcribed using Sp6 mMessage mMachine kit (Ambion). Using a micro-injector, 100 pg/embryo of *sox32* mRNA was injected in donor homozygous *Tg(ins:DsRed)* embryos. Embryos from an *ins +/-* incross served as host embryos. Between the 1k-cell and sphere stage (3-4 hpf), 15-20 cells from the donor were transplanted to host embryos, targeting the host mesendoderm, at the margin of the blastoderm.

### Wholemount in situ hybridization

Larvae were collected at 120 hpf and fixed with 4% paraformaldehyde in PBS overnight at 4 °C. In situ hybridization was performed as described previously (Thisse & Thisse, 2008). *ar* Digoxigenin-labelled anti-sense probe was synthesized using T7 polymerase (Roche) and DIG RNA labelling kit (Roche). The probe template was amplified using the following primers: *ar* ISH-forward 5’-TGGAGTTTTTCCTTCCTCCA-3’ and *ar* ISH-reverse 5’-TAATACGACTCACTATAGGGTCATTTGTGGAACAGGATT- 3’, obtaining a 1100 bp probe as described previously (Gorelick et al., 2008). Embryos were imaged on a Nikon SMZ25 stereomicroscope. Mutant and wild-type larvae were processed in the same tube and genotyped after the images were taken.

### Drug screening

At 72 hpf, 3 *ins* mutant larvae were pooled in each well of a 96-well plate in 200 μl of egg water buffered with 10mM HEPES. All drug treatments were performed at 10 μM with 1% DMSO, unless otherwise stated. Drug treatment was performed from 84 hpf to 120 hpf, after which each well was visually analyzed to assess toxicity. Subsequently, 100 μl of egg water was removed and 25 μl of 5X Cell Culture Lysis Buffer (Promega) was added. The plate was left shaking for 1 min at 750 rpm, and after gentle shaking at 150 rpm for 30 min, another round of vigorous shaking was performed for 1 min at 750 rpm. 8 μl from each well was used for the glucose detection reaction in a new 96-well plate using the Glucose Assay Kit (CBA086, Merck). All drug treatments were performed at 10 μM, unless indicated otherwise. Drugs with an average glucose-lowering effect of over 10% were retested.

### Drug libraries used in this screen are

a. 1440 molecules from Edelris Keymical Collection (Edelris) (0 hits).
b. 285 molecules from the CLOUD collection (Licciardello et al., 2017) (2 hits).
c. 156 molecules identified as *insulin* stimulators (Matsuda et al., 2018) (1 hit).
d. 352 kinase inhibitors (SelleckChem) (0 hits)

### Transcriptomic analyses

For RNA-seq analysis, total RNA was isolated from 120 hpf zebrafish using the RNA Clean & Concentrator kit (Zymo Research) combined with DNase digestion (RNase-free DNase Set, Promega) to avoid contamination by genomic DNA. RNA and library preparation integrity were verified with LabChip Gx Touch 24 (Perkin Elmer). 3μg of total RNA was used as input for Truseq Stranded mRNA Library preparation following manufacture’s low sample protocol (Illumina). Sequencing was performed on NextSeq500 instrument (Illumina) using v2 chemistry, resulting in minimum of 23M reads per library with 1×75bp single end setup. The resulting raw reads were assessed for quality, adapter content and duplication rates with FastQC (Andrews, 2010). Trimmomatic version 0.33 was employed to trim reads after a quality drop below a mean of Q20 in a window of 5 nucleotides (Bolger et al., 2014). Only reads between 30 and 150 nucleotides were cleared for further analyses. Trimmed and filtered reads were aligned versus the Ensembl Zebrafish genome version DanRer10 (GRCz10.90) using STAR 2.4.0a with the parameter “-outFilterMismatchNoverLmax 0.1” to increase the maximum ratio of mismatches to mapped length to 10% (Dobin et al., 2013). The number of reads aligning to genes was counted with featureCounts 1.4.5-p1 tool from the Subread package (Liao et al., 2014). Only reads mapping at least partially inside exons were admitted and aggregated per gene. Reads overlapping multiple genes or aligning to multiple regions were excluded. Differentially expressed genes were identified using DESeq2 version 1.62 (Love et al., 2014). Maximum Benjamini-Hochberg corrected p-value of 0.05, and a minimum combined mean of 5 reads were set as inclusion criteria. The Ensemble annotation was enriched with UniProt data (release 06.06.2014) based on Ensembl gene identifiers (“Activities at the Universal Protein Resource (UniProt),” 2014). RNA-seq data have been deposited in the ArrayExpress database at EMBL-EBI (www.ebi.ac.uk/arrayexpress) under accession number <u>E-MTAB-7283</u>.

For microarray expression profiling, RNA was isolated from pooled 108 hpf zebrafish larvae using the RNA Clean & Concentrator kit (Zymo Research) combined with DNase digestion (RNase-free DNase Set, Promega). 10 animals were used for each pooled sample. Sample quality was tested using a Bioanalyzer and microarray analysis was performed by Oak Labs (Germany). Microarray data have been deposited in the ArrayExpress database at EMBL-EBI (www.ebi.ac.uk/arrayexpress) under accession number <u>E-MTAB-7282</u>.

### Mass spectrometric analysis

For each of the three biological replicates within a genotype, protein was extracted from pools of 600 larvae at 5 dpf using 4% SDS in 0.1 M Tris/HCl, pH 7.6 and a tissue disrupting sterile pestle (Axygen) for lysis. After heating to 70°C at 800 rpm for 10 min and DNA shearing by sonication, cell debris was removed by centrifugation at 14.000 × g for 10 min and retaining the supernatant. Using DC protein assay (BioRad),7 mg of solubilized proteins per sample were Acetone precipitated at -20°C, overnight, followed by centrifugation at 14.000 × g for 10 min and washing the pellet washing using 90% acetone. After evaporation of residual acetone, samples were dissolved in urea buffer (6 M urea, 2 M thiourea, 10 mM HEPES, pH 8.0), followed by enzymatic peptidolysis as described (Graumann et al., 2008) with the following adaptations: 10 mM dithiothreitol, 55 mM iodoacetamide and 100:1 protein to enzyme ratio of the proteolytic enzymes were used. The resulting peptide concentration was estimated using the Fluorimetric Peptide Assay (Pierce) and peptides were labelled by reductive demethylation as described (Boersema et al., 2009) using light (L) and heavy (H) labels for wild type and mutants respectively. Differentially labelled peptides were mixed at 1:1 and subjected to a high pH reversed-phase peptide fractionation kit (Pierce). Full proteome analysis from the resulting eight fractions was performed using an EASY-nLC 1000 UHPLC system (ThermoFisher Scientific) and 20 cm in-house packed C_18_ capillary emitter columns (70 μm inner column diameter, 1.9 μm C18 beads, Dr. Maisch GmbH) coupled to a QExactive HF orbitrap mass spectrometer (ThermoFisher Scientific) using an electrospray ionization source. The gradient employed consisted of linearly increasing concentrations of solvent B (90% acetonitrile, 1% formic acid) over solvent A (5% acetonitrile, 1% formic acid) from 5% to 30% over 215 min, from 30% to 60%, from 60% to 95% and from 95% to 5% for 5 min each, followed by re-equilibration with 5% of solvent B. The constant flow rate was set to 400 nl/min.

Full MS spectra were collected for a mass range of 300 to 1750 m/z with a resolution of 60,000 at 200 m/z. The ion injection target was set to 3 × 10^6^ and the maximum injection time limited to 20 ms. Ions were fragmented by higher energy collision dissociation (HCD) using a normalized collision energy of 27, an isolation window width of 2.2 m/z and an ion injection target of 5 × 10^5^ with a maximum injection time of 20 ms. Precursors characterized with unassigned charge state and a charge state of 1 were excluded from selection for fragmentation. Precursor resequencing was prevented using dynamic exclusion time of 20 s. Resulting tandem mass spectra (MS/MS) were acquired with a resolution of 15,000 in a top 15 loop.

Data analysis: MS raw data were processed by MaxQuant (1.6.0.1) (Cox & Mann, 2008) using the Uniprot zebrafish database (as of 20.04.2017) containing 59066 entries and the following parameters: a maximum of two missed cleavages, mass tolerance of 4.5 ppm for the main search, trypsin as the digesting enzyme, carbamidomethylation of cysteines as a fixed modification and oxidation of methionine as well as acetylation of the protein N-terminus as variable modifications. For the dimethyl-labeled protein quantification, isotope labels were configured for peptide N-termini and lysine residues with a monoisotopic mass increase of 28.0313 and 36.0757 Da for the light and heavy labels, respectively. Peptides with a minimum of seven amino acids were included in the analysis. MaxQuant was set to filter for 1% false discovery rate on the peptide and protein levels, both. Only proteins with at least two peptides and one unique peptide were considered identified and included in further data analysis.

Canonical Pathway Analysis was performed using Ingenuity Pathway Analysis (IPA) (Qiagen). Differentially expressed proteins from our study (log2FC+-1.5) and from previously published datasets were subjected to a Comparison Analysis in IPA. P-value maximum cut-off was set at 0.05 and the processes are listed according to those affected across the most studies.

## Acknowledgements

We would like to thank Dr. Rashmi Priya and Dr. Sonja Sievers for discussion and comments, Dr. Radhan Ramadass for microscopy, Martin Laszczyk and the rest of the animal caretaking team for zebrafish care. These studies were supported by funding from the Max Planck Society (DYRS).

## Ethics Statement

Zebrafish husbandry was performed under standard conditions in accordance with institutional (MPG) and national ethical and animal welfare guidelines approved by the ethics committee for animal experiments at the Regierungspräsidium Darmstadt, Germany.

---

**Figure 1 - figure supplement 1.**
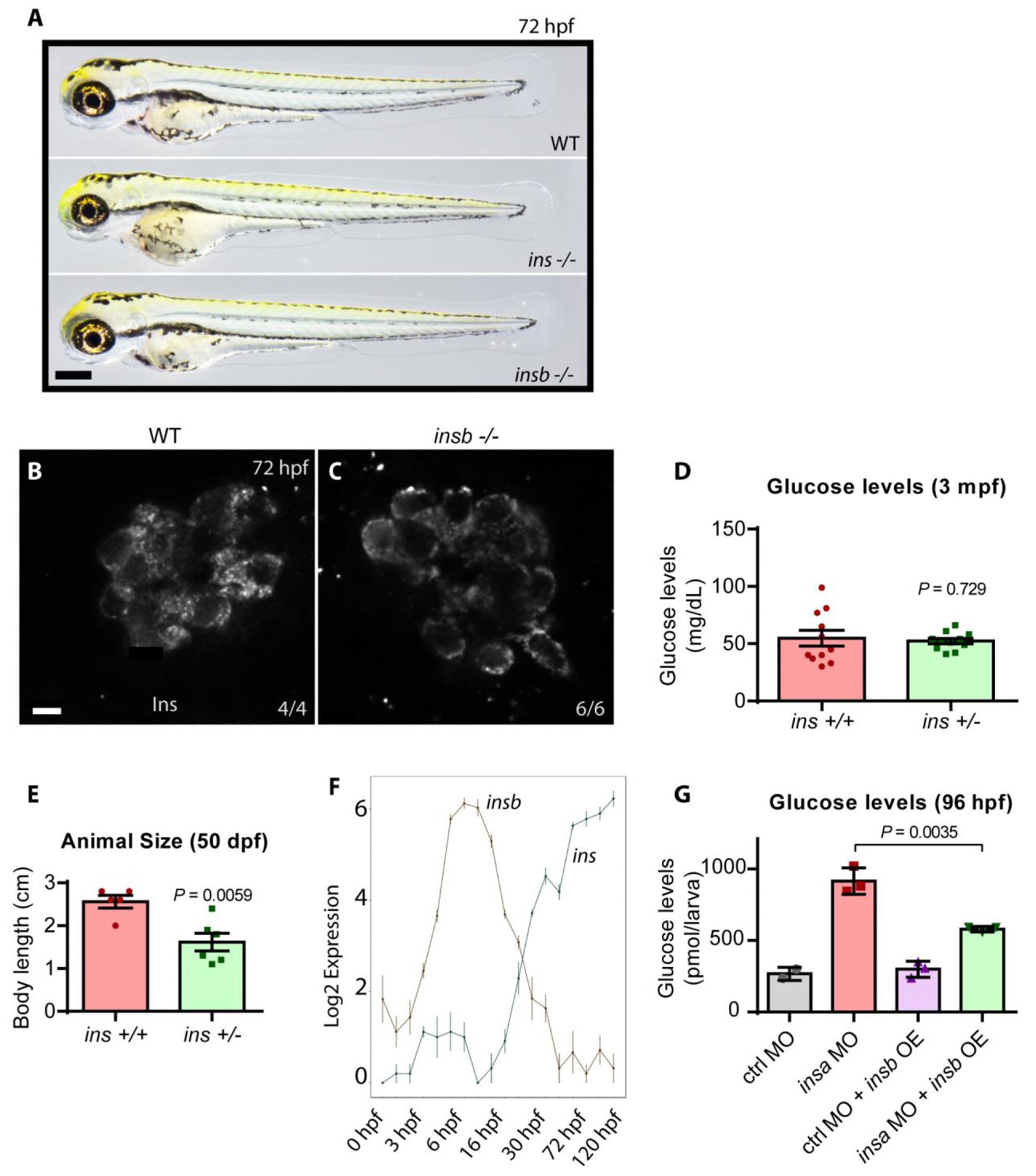
*ins*, and not *insb*, is the predominant paralog expressed in zebrafish pancreatic islets. **A)** Brightfield images of 72 hpf wild-type, *ins* mutant and *insb* mutant larvae. B-C) Confocal planes of pancreatic islets in 72 hpf wild-type and *insb* mutant larvae stained for Insulin (white). **D)** Blood glucose levels from wild-type and *ins +/-* 3 mpf adult animals; mean ± SEM, n=11 animals. **E)** Body length measurements from wild-type and *ins +/-* 50 dpf juvenile animals; mean ± SEM, n=5-6 animals. **F)** mRNA expression time course of *ins* and *insb* transcripts during zebrafish development from 0 to 120 hpf. **G)** 4 ng of control (ctrl) or *ins* morpholino (MO) was injected in non-transgenic or *Tg(ins:insb)* expressing *(insb* OE) embryos at the 1-cell stage and glucose levels were measured at 96 hpf; mean ± SEM, n=3 replicates. Scale bars: 250 μm (A), 5 μm (C, **D)**.

**Figure 2 - figure supplement 1.**
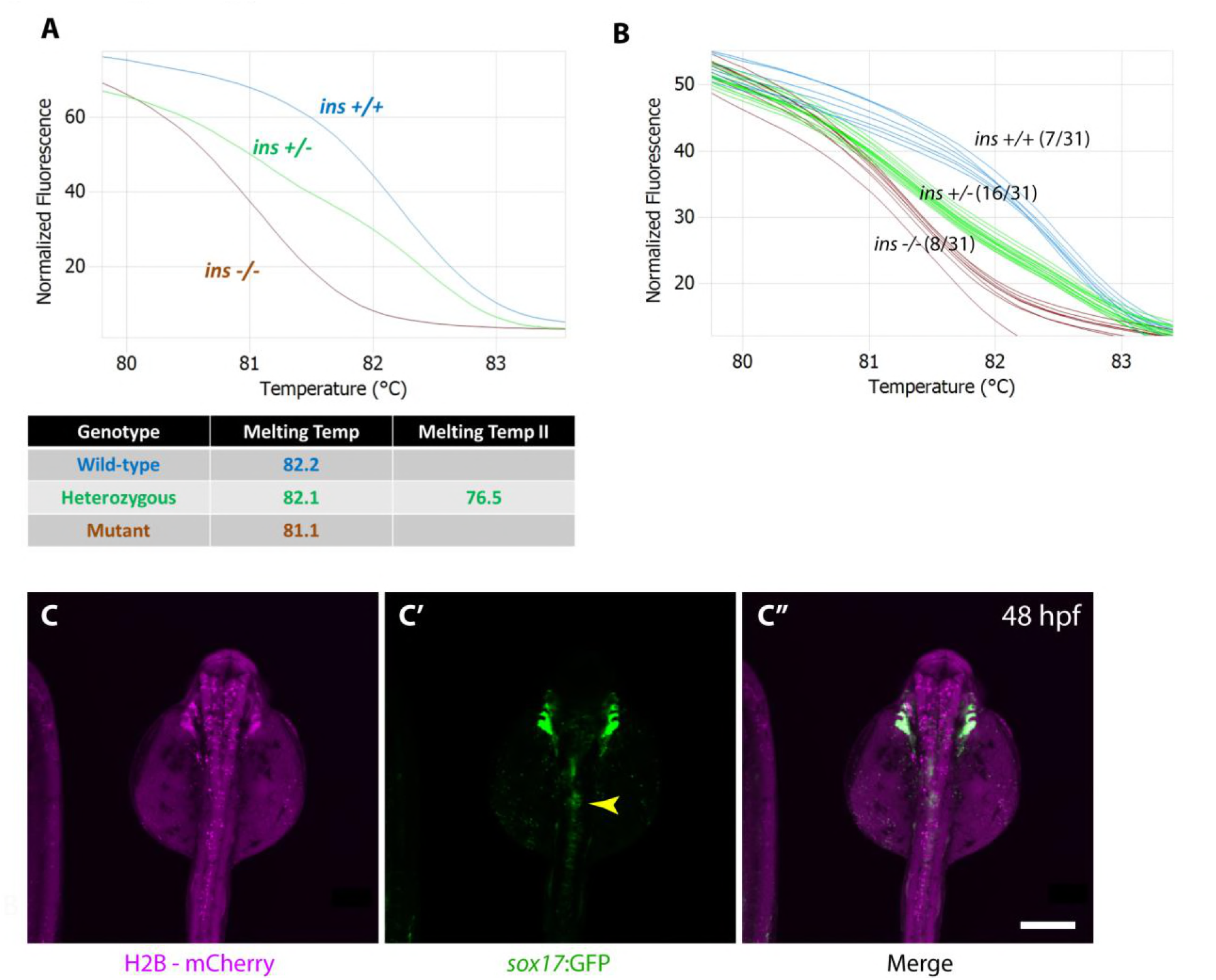
*sox32* mRNA-injected cells contribute to host endoderm upon transplantation. **A)** High-resolution melt analysis patterns for genotyping *ins +/+* (blue), *ins +/-* (green) and *ins -/-* (brown) animals. **B)** Representative example of genotyping 31 chimeric adults from fin tissue, revealing 8 of 31 animals to be mutant (brown). C-C”) Confocal images of a 48 hpf host embryo injected with *H2B-mCherry* mRNA (magenta) and transplanted with donor *Tg*(*sox17*:GFP) expressing endoderm showing the chimeric islet (yellow arrowhead). Scale bar: 200 μm.

**Figure 3 - figure supplement 1.**
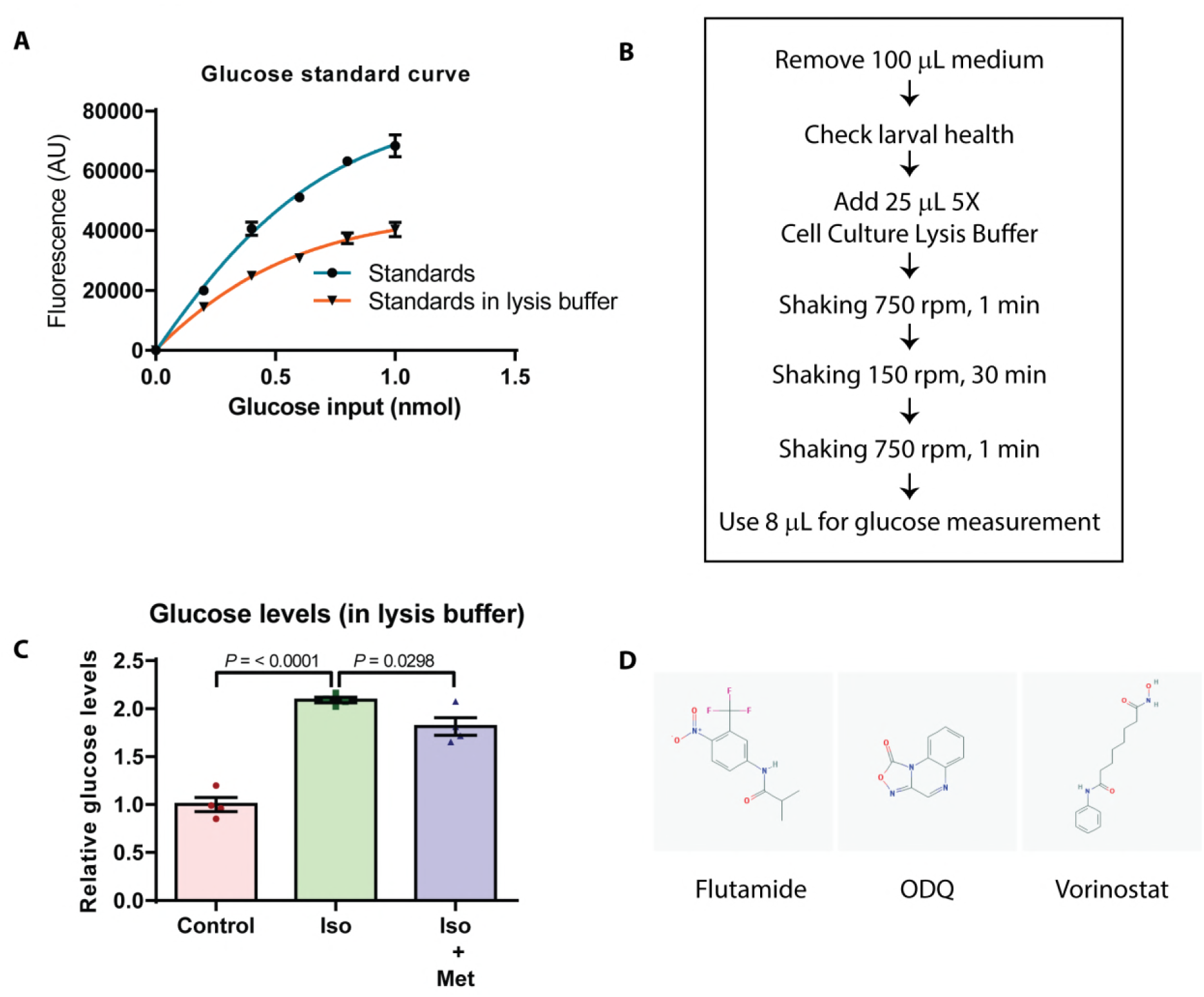
A 96-well plate-adapted protocol to measure glucose levels is suitable for small molecule screening. **A)** Plotting the measured glucose interpolated from the regular standard curve (blue) shows a shifted curve in the presence of lysis buffer (orange) with a dynamic range still suitable for measuring glucose. **B)** Schematic showing the plate-adapted glucose readout assay. **C)** Relative glucose levels upon treatment of wild-type larvae from 4 to 6 dpf with beta-adrenergic agonist Isoprenaline (Iso) or both Isoprenaline and 250μM Metformin (Iso + Met); mean ± SEM, n = 4-5 replicates. **D)** Structures of the hit molecules - Flutamide, ODQ and Vorinostat.

**Figure 4 - figure supplement 1.**
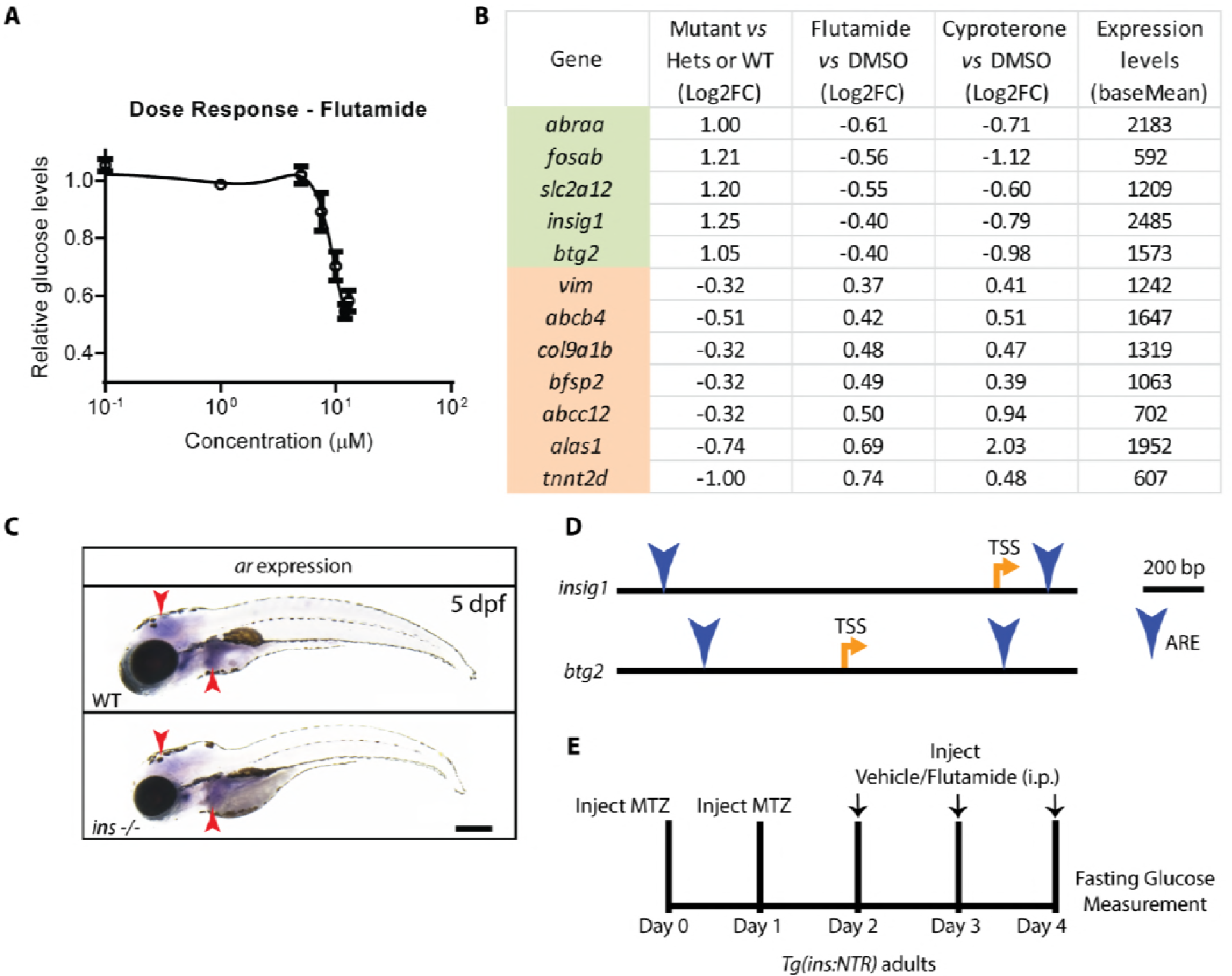
Flutamide reduces glucose in a dose-dependent manner, possibly exerting its effects through liver gluconeogenic enzymes. **A)** Dose response curve of the glucose lowering effect of Flutamide on *ins* mutant larvae, treated from 84 to 120 hpf; mean ± SEM, n = 3 replicates. **B)** The 12 genes differentially regulated in *ins* mutants compared to non-mutant siblings and modulated in the opposite direction upon treatment with Flutamide or Cyproterone, compared to DMSO; listed with their fold change (Log_2_ scale) and expression levels under control condition (base mean); genes are ordered by the fold change under Flutamide vs DMSO condition. **C)** Wholemount in situ hybridization for *ar* transcripts in 120 hpf wild-type and *ins* mutant larvae showing expression in the brain and liver (red arrowheads). **D)** Schematic of *insig1* and *btg2* gene loci, with locations of Transcription Start Site (TSS) and Androgen Response Elements (ARE, blue arrowheads) indicated. **E)** Schematic of vehicle vs Flutamide treatment of *Tg(ins:NTR)* adult animals following J3-cell ablation with Metronidazole (MTZ) injection to induce hyperglycemia. Scale bar: 250

## Titles for Supplementary Files

**Supplementary file 1:** List of proteins with Log2FC > 1 or Log2FC < -1 from proteomic analyses comparing 120 hpf *ins* mutant and wild-type animals.

